# Shiitake mycelium fermentation improves digestibility, nutritional value, flavor and functionality of plant proteins

**DOI:** 10.1101/2021.10.07.463529

**Authors:** Anthony Clark, Bhupendra K Soni, Brendan Sharkey, Terry Acree, Edward Lavin, Hannah M. Bailey, Hans H. Stein, Ashley Han, Marc Elie, Marina Nadal

## Abstract

Plant proteins can serve as inexpensive and environmentally friendly meat-replacements. However, poor taste characteristics and relatively low nutritional value prevent their full acceptance as meat substitutes. Fermentation of food has been historically used to improve the quality of foods. In this work we describe the improvement in digestibility, nutritional value, physical properties, and organoleptic characteristics, of a pea and rice protein concentrate blend through fermentation with shiitake mushroom mycelium. Ileal digestibility pig studies show increases in the DIAAS for the shiitake fermented pea and rice protein blend turning the blend into an “excellent source” of protein for humans. The fermentation also increases the solubility of the protein blend and reduces the content of the antinutrient compounds phytates and protease inhibitor. Mass spectrometry and sensory analyses of fermented protein blend indicates that fermentation leads to a reduction in off-note compounds substantially improving its organoleptic performance.

## INTRODUCTION

Plant-based protein foods are emerging as alternative to animal derived protein (Sexton et al., 2019). Several advantages make plant protein an ideal replacement to meat; however, two main drawbacks prevent their full acceptance in the food space. In general, the nutritional value of unprocessed single source plant protein for humans is often inferior to that of animal protein sources. By themselves, proteins derived from pea (*Pisum sativum*) and rice (*Oryza sativa*) are deficient in lysine, methionine and some branched-chain amino acids (Gorissen et al., 2018), and are therefore considered of lower nutritional quality (USDA, 2013). However, if combined in correct proportions, pea protein and rice protein may complement each other to deliver a blend with an ideal balance of indispensable amino acids that is adequate for human nutrition. In 1991 the Food and Agriculture organization (FAO) and World Health Organization (WHO) introduced the Protein Digestibility Corrected Amino Acid Score (PDCAAS)(FAO/WHO, 1991). This concept is based on the assumption that a protein blend’s nutritional value is determined not only by the amino acid profile, but also by the ability of the human gastrointestinal tract to hydrolyze individual proteins and by the rate at which free amino acids are absorbed into the blood stream (FAO, 2013). Although the PDCAAS score has been widely adopted to describe protein nutritional value, it is calculated from the total tract digestibility of crude protein (CP) and based on the assumption that all amino acids (AAs) in CP have the same digestibility. However, the digestibility of CP is not representative of the digestibility of all AAs because individual AAs are digested with different efficiencies (Stein et al., 2007). Moreover, fermentation of the free AAs by the lower intestine microbiome can affect fecal AA excretion and hence alter the PDCAAS values (Sauer & Ozimek, 1986). Therefore, measuring digestibility at the distal ileum (the end of the small intestine) provides the most realistic estimate of AA bioavailability as compared to total tract digestibility (Cervantes-Pahm et al., 2014). Based on these facts, in 2013 the FAO introduced the Digestible Indispensable Amino Acid Score (DIAAS) as a method to evaluate protein quality (Wolfe et al., 2016). Because DIAAS is calculated by measuring ileal digestibility of individual AAs, it more accurately describes the true nutritional value of dietary protein than the PDCAAS method (Bailey & Stein, 2019). Additionally, DIAAS method provides a more precise assessment of protein quality for a blend of different dietary protein sources. Nonetheless, PDCAAS is still widely used in North America as measurement of protein quality.

Protein digestibility is also partially dependent on the solubility of the protein material and the presence of residual antinutrients such as protease inhibitors and phytic acid (Afify et al., 2012). Cereal grains and legumes contain several protease inhibitors of major concern (Samtiya et al., 2020). Particularly pea is rich in trypsin inhibitors (Avilés-Gaxiola et al., 2018) while rice bran is known to contain considerable amounts of the oryzacystatin-I (OC-I), a rice cystatin (cysteine protease inhibitor) which binds tightly and reversibly to the papain-like group of cysteine proteinases (Abe et al., 1987). Although mounting scientific evidence is starting to reveal extended health benefits of plant antinutrients (Lajolo & Genovese, 2002), the removal/reduction of such compounds in plant protein concentrates remains highly desirable to increase digestibility of proteins. More often, antinutrients complex with proteins forming precipitates that are not easily accessible by gastric digestive enzymes (Joye, 2019). Phytic acid is the main storage of phosphorous in seeds of legumes and cereals (Reddy et al., 1982). Due to its 6 phosphate groups, phytic acid acts as a powerful chelating agent, interfering with absorption of key minerals such as zinc, iron, magnesium and calcium in the gastrointestinal tract during digestion (Bohn et al., 2008). Moreover, because phytate can sequester Ca^2+^ and Mg^2+^, co-factors of digestive proteases and α-amylases, it can indirectly impair digestion (Deshpande & Cheryan, 1984; Khan & Ghosh, 2013). A direct inhibitory effect of phytate on these enzymes has also been proposed (Sharma et al., 1978). Therefore, the presence of phytate in protein concentrates has the potential of negatively impacting digestibility in several ways and consequently lowering the nutritional quality of plant proteins. Removal of phytates greatly improves the nutritional value of foods and several methodologies are employed in the food industry to eliminate their presence (Gupta et al., 2015). Phytases, the enzymes responsible for hydrolyzing phytic acid into inositol and phosphate (Lei et al., 2013) are widely distributed among microorganisms, including fungi such as shiitake (Jatuwong et al., 2020).

The other main disadvantage of plant derived proteins are their undesirable organoleptic characteristics. Specifically, plant proteins often display off-flavors, which makes their incorporation into meat or dairy analog products challenging. For example, plant proteins such as pea proteins are associated with beany aromas due to the presence of the volatiles 3-alkyl-2-methoxypyrazines (galbazine) and have bitter flavors associated with plant lipids and saponins (Gläser et al., 2020; Roland et al., 2017).

In this work we describe the improvement in digestibility, nutritional value, and organoleptic characteristics of FermentIQ^®^ protein (Soni Bhupendra, Kelly Brooks, Langan Jim, Hahn Alan, 2018), a shiitake-fermented pea-rice protein concentrate blend, as compared to the same unfermented pea and rice protein blend.

## MATERIALS AND METHODS

### Ileal Digestibility Studies

Two diets were formulated with the unfermented and fermented protein blends included in one diet each as the only AA containing ingredient. The third diet was a nitrogen-free diet that was used to measure basal endogenous losses of CP and AA. Vitamins and minerals were included in all diets to meet or exceed current requirement estimates for growing pigs. All diets also contained 0.4% titanium dioxide as an indigestible marker, and all diets were provided in meal form.

Nine castrated male pigs at 10 weeks of age (initial BW: 28.5 ± 2.3 kg) were equipped with a T-cannula in the distal ileum and allotted to a triplicated 3 × 3 Latin square design with 3 pigs and 3 periods in each square. The number of pigs exceeded the recommended minimum number of pigs needed to obtain reliable values for DIAAS(FAO, 2014). Diets were randomly assigned to pigs in such a way that within each square, one pig received each diet, and no pig received the same diet twice during the experiment. Therefore, there were 9 replicate pigs per treatment. Pigs were housed in individual pens (1.2 x 1.5 m) in an environmentally controlled room. Pens had smooth sides and fully slatted tribar floors. A feeder and a nipple drinker were also installed in each pen.

All pigs were fed their assigned diet in a daily amount of 3.3 times the estimated energy requirement for maintenance (i.e., 197 kcal ME per kg^0.60^). Feeding and collection of fecal samples and ileal digesta samples followed procedures described previously (Mathai et al., 2017).

At the conclusion of the experiment, ileal samples were thawed, mixed within animal diet, and a sub-sample was collected for chemical analysis. Ileal digesta samples were lyophilized and finely ground prior to chemical analysis. Fecal samples were dried in a forced-air oven and ground through a 1 mm screen in a Wiley Mill (model 4, Thomas Scientific) prior to chemical analysis. All samples were analyzed for dry matter (DM; Method 927.05) and for CP by combustion (Method 990.03) at the Monogastric Nutrition Laboratory at the University of Illinois Champagne, IL. All diets, fecal samples, and ileal digesta were analyzed in duplicate for titanium (Method 990.08; Myers et al., 2004). The two proteins, all diets, and ileal digesta samples were also be analyzed for AA [Method 982.30 E (a, b, c)](Horowitz et al., 1957).

Values for apparent ileal digestibility (AID) and standardized ileal digestibility (SID) of CP and AA were calculated (Stein et al., 2007), and standardized total tract digestibility (STTD) of CP were calculated as well (Mathai et al., 2017). Values for STTD and SID were used to calculate values for PDCAAS and PDCAAS-like, and DIAAS, respectively, as previously explained (Leser, 2013; Mathai et al., 2017).

The protocol for the animal work was reviewed and approved by the Institutional Animal Care and Use Committee at the University of Illinois (Protocol Number 16113).

#### Solubility

Solubility of protein samples was calculated as:

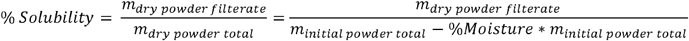

Sample moisture was calculated after placing 5g of protein powder in a desiccator and recording the dried weight, as follows: 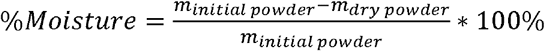

Dry powder filtrate was calculated by dissolving 2.5 g of dried sample in 50 ml at room temperature and adjusting the pH to either 3, 5, 6, 7, or 8, with 1M HCl or 1M NaOH. Samples were mixed thoroughly and centrifugated at 9000 RPM for 10 minutes. Supernatant was vacuum filtrated using GE Whatman 47mm Grade 4 filter papers (GE) and the weight recorded.

#### Phytate Measurement

Phytic acid was measured by Eurofins Scientific, Luxembourg, by the method of stable phytate-iron complex formation in dilute acid solution.

#### Enzyme inhibition assays

Trypsin inhibition assay was performed by Eurofins Scientific (Method AOC S Ba 12-75). Chymotrypsin was performed by Reaction Biology Corporation, Malvern, PA (https://www.reactionbiology.com). Papain and subtilisin inhibition assays were performed as previously described ^43^. Briefly, inhibitory activity was assessed by incubating 0.5 mL extract of fermented product with 0.5 mL of commercial papain (EC 3.4.22.2) or subtilisin (EC 3.4.21.62) and incubating at 37°C for 15 min. Then, 5 mL of a casein solution (0.65% w/v) was added to the assay solution and the mixture was further incubated at 55°C for exactly 10 min. Inhibitory activity was measured by obtaining the difference between the enzyme activity in the absence and in the presence of the fermented protein blend.

#### GC-O and CHARM (Combined Hedonic Aroma Response Measurement) analysis

identification of volatile compounds in fermented and unfermented protein blend samples was done by gas chromatography/olfactometry (GC/O) using human “sniffers” to assay for odor activity among volatile analytes as previously described (Acree & van Ruth, 2003).

#### Sensory Panel Assessment

The powdered unfermented and fermented protein blend samples were used at 10% in room temperature water and mixed. Sensory testing was performed by Sensation Research, Mason, OH (https://sensationresearch.com/) using a combination of Spectrum Method™ and Quantitative Descriptive Analysis (QDA) (Hootman et al., 1992). Trained descriptive panelists used full descriptive analysis technique to develop the language, ballot, and rate profiles of the products on aroma. Eleven panelists were trained for 2 sessions with 2 individual evaluations per sample for data collection. Eleven trained panelists (experienced from prior protein consensus panels) evaluated appearance for all samples immediately after mixing to capture initial scores and minimize variability. Data were analyzed using Senpaq: Descriptive Analysis - Analysis of variance (ANOVA).

## RESULTS

### Digestibility of fermented pea and rice protein blend

The use of pigs as models for humans was recommended due to the impracticality of obtaining ileal digesta from humans and because pigs are better models for humans than rats (FAO, 2013). Subsequently, DIAAS in both animal and plant proteins have been determined using the pig model the same way as was done in this experiment (Cervantes-Pahm et al., 2014; Mathai et al., 2017)

Nutritional analysis of the unfermented and fermented protein blends indicated that the CP content was similar in both samples with 77.57% and 76.77%, respectively (Table 1). Concentrations of total indispensable AA were also similar in the two protein blends, with the unfermented blend containing 37.51% and the fermented blend containing 35.88%. However, the concentration of Lysine (Lys) was approximately 25% greater in the unfermented sample compared with the fermented sample.

The Apparent Ileal Digestibility (AID) and Standardized Ileal Digestibility (SID) of CP did not differ between the unfermented and the fermented pea-rice protein concentrate (Table 2). In addition, the AID and SID of all indispensable and dispensable AA did not differ between the two proteins.

The PDCAAS values were calculated using the FAO recommended scoring patterns(Leser, 2013) for “young children” (6 months to 3 years) and for “older children, adolescents, and adults” (3+ years) (Table 3), and found not to be different between unfermented and fermented protein blends for both age groups. For young children, PDCAAS values were similar to those calculated for children 2 to 5 years, with the unfermented and fermented proteins having PDCAAS values of 86 and 91, respectively. For PDCAAS values calculated for older children, unfermented and fermented proteins had values of 101 and 108, respectively. The first limiting AA when compared with the AA requirements was SAA and Lys for unfermented protein and fermented protein, respectively, for both age groups.

DIAAS was calculated for “young children” and for “older children, adolescents, and adults” (Sotak-Peper et al., 2017) (Table 4). The DIAAS values calculated for both age groups were greater (*P* < 0.05) for the fermented than for the unfermented pea-rice protein. For young children, the DIAAS was 70 and 86 for unfermented and fermented proteins, respectively, which represents a 23% increase. For older children, adolescents, and adults, the DIAAS was 82 and 102 for unfermented and fermented proteins, respectively, which represents a 24% increase. The first limiting AA in the proteins when compared with the AA requirements for both age groups was SAA and Lys for unfermented and fermented proteins, respectively.

### Solubility and antinutrient levels of fermented pea and rice protein blend

To determine if the fermentation process also impacts physical properties of the pea and rice protein blend, the solubility of the fermented and unfermented protein concentrate blends was calculated across a wide range of pH. The dissolved solids of three independently fermented protein blend samples were consistently higher than that of unfermented protein blend (raw pea + rice) showing an increase at all pH values (Figure 1). The minimal increase in dissolved solids in the fermented samples over the mixture of raw materials was 2-fold and occurred at pH 5, while the highest increase was 3-fold, at pH 8.

**Figure 1.**
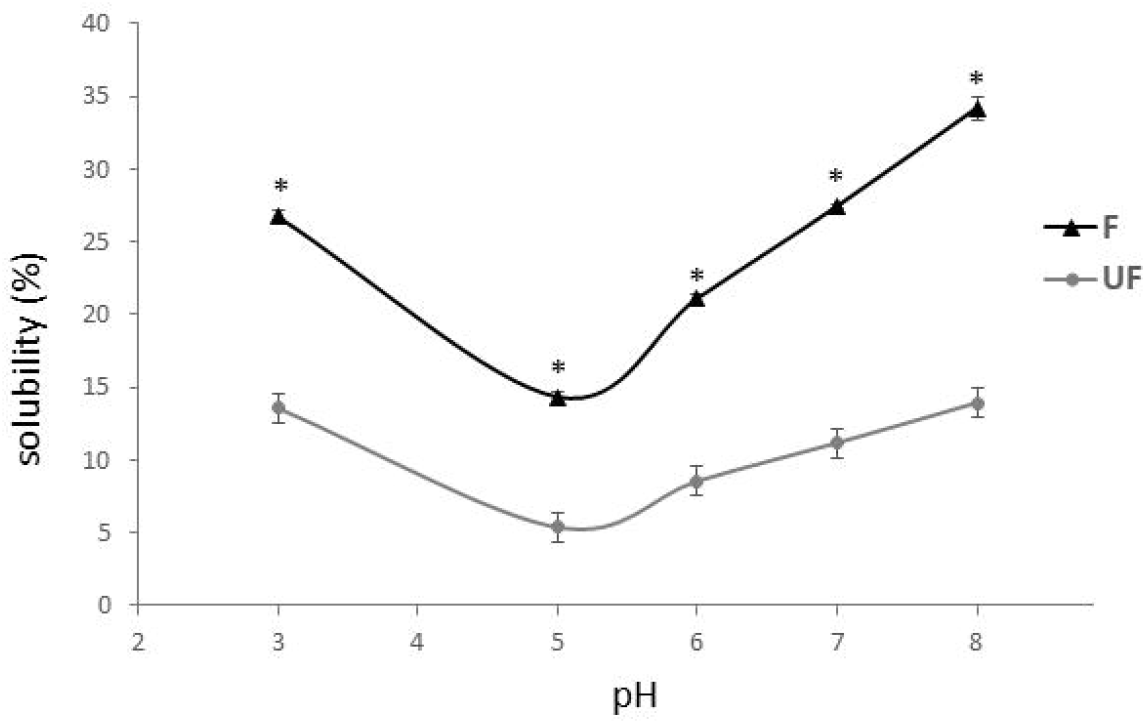
Changes in solubility during with fermentation process. The solubility of unfermented (**UF**) and fermented (**F**) protein blend was evaluated at pH 3, 4, 5, 6, 7 and 8. Values represent the main of 3 technical replicates. Error bars express standard error. Asterisks indicate significant difference (t-test; P < 0.01).

To assess the reduction of protein inhibitors of key proteases due to the fermentation process, inhibitory enzyme assays were conducted. No changes in trypsin, chymotrypsin and subtilisin inhibition were observed between unfermented and fermented protein blends (data not shown). A substantial reduction in papain inhibition was observed when comparing unfermented (3.4 IU/g protein bled) in comparison to the unfermented (0.6 IU/g protein protein) blend (Figure 2).

**Figure 2.**
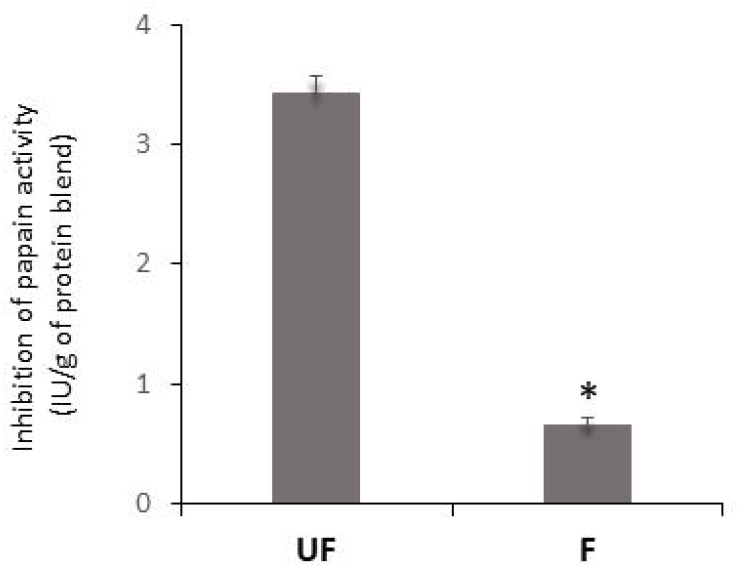
Quantification of papain activity. Papain enzyme inhibition was evaluated in the presence of unfermented (UF) and fermented (F) protein blends. Asterisks indicate significant difference (t-test; P <0.01).

The presence of residual phytate in plant protein can negatively affect protein digestibility. To evaluate the ability of shiitake fermentation to remove phytic acid the levels of phytate were measured in both unfermented and fermented protein blends. The percentage of phytic acids in the unfermented and fermented protein blends were 1.25% and 0.68%, respectively. These results indicate changes in physical properties and chemical composition of the fermented protein blend.

### Organoleptic characteristics of fermented pea and rice protein blend

To characterize and quantify changes in volatile compounds associated with the organoleptic profile of unfermented and fermented pea and rice protein concentrate mixtures, both protein blends were subjected to GC-MS and GC-olfactometry and Combined Hedonic Aroma Response Measurement (CHARM) analyses. The results indicate a decrease in the earthy, beany, potato and mustard off-notes in the fermented protein blend compared to the unfermented, while those associated with fatty and musty are increased (Figure 3, Supp Table1-5). The analysis also indicates an overall change in the relative abundance of volatile compounds in the fermented protein blend as compared to the unfermented one (Figure 4A). Several compounds, including galbazine, methyl mercaptan, methional and a sesquiterpene similar to bergamotene (bergamotene-like) were described as imparting unpleasant off-flavors. Specifically, off-notes compounds methional, methyl mercaptan, bergamotene-like compound which are present in the unfermented protein blend were substantially reduced in the fermented protein blend by 40%, 78%, 99% respectively. Moreover, the potent beany off-notes associated with (galbazine) present in the unfermented protein blend were not detected in the fermented sample (Figure 4B). To further understand the aroma profile of the fermented and unfermented protein blends, a sensory evaluation was carried out by a trained sensory panel of 11 eleven people. The sensory results correlate well with data from CHARM analysis, indicating a statistically significant decrease in pea and rice notes and overall improvement aroma of the fermented blend (Table 5). The GC-MS data also reveals a relative increase in the oxylipins: 1-octen-3-one; 2,6-decadienal; 2,4-nonadienal and 2,3 butanedione in the fermented protein blend as compared to the unfermented blend (Figure 4A; Supp Table 1; Suppl Figure1), however this change was not reflected in the sensory profiles provided by the sensory panel. In fact, 2,3 butanedione had a positive impact to the sensory profiling of the fermented protein blend. All together, these results indicate an improvement in the organoleptic characteristic in the fermented pea and rice protein concentration blend versus the unfermented protein blend.

**Figure 3.**
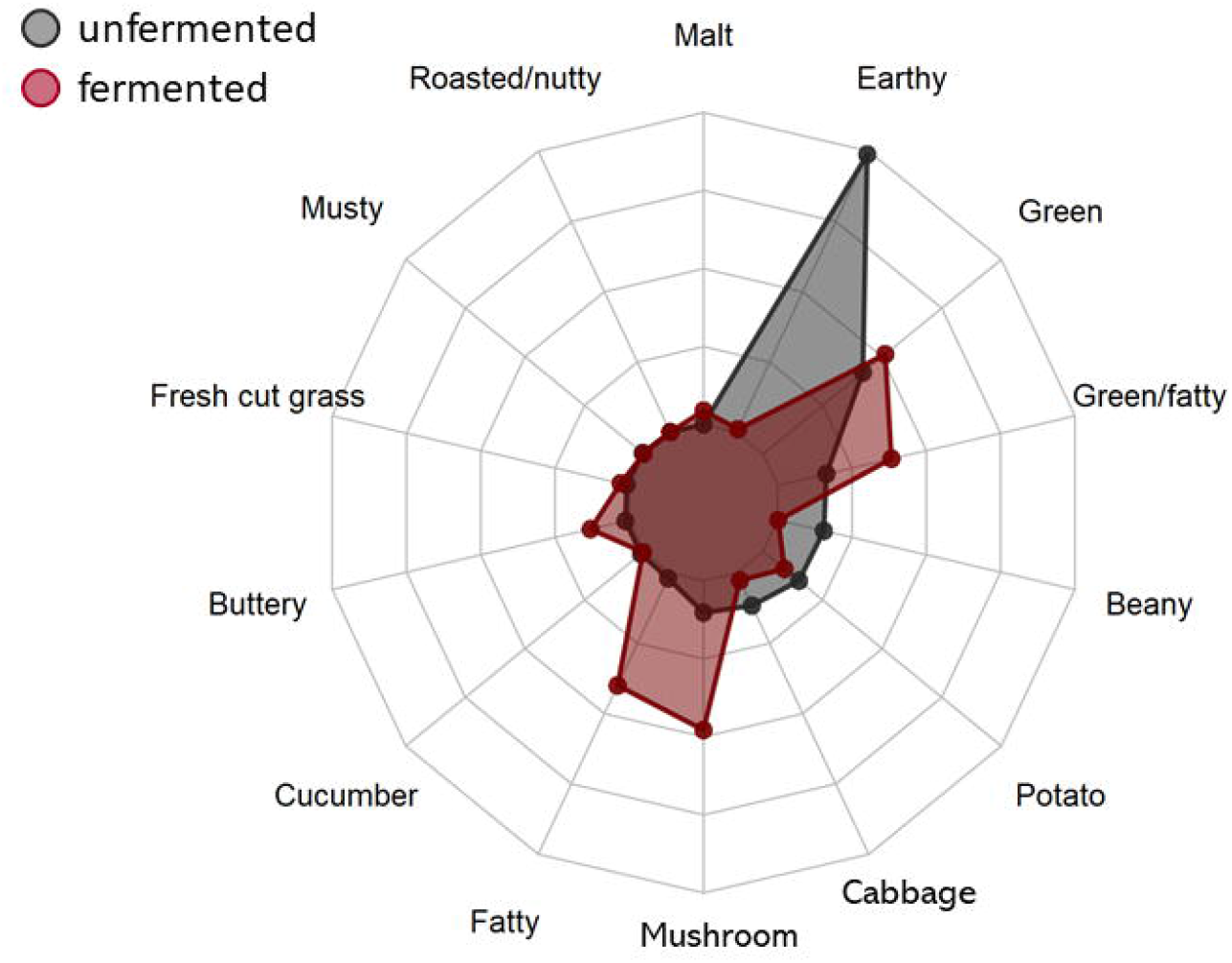
Odorant profile analysis of fermented and unfermented protein blends. GC-olfactometry and Combined Hedonic Aroma Response Measurement (CHARM) analyses of fermented and unfermented protein blends. Only attributes that are significantly different at the 90% confidence level as tested by ANOVA are shown in the spider plot.

**Figure 4.**
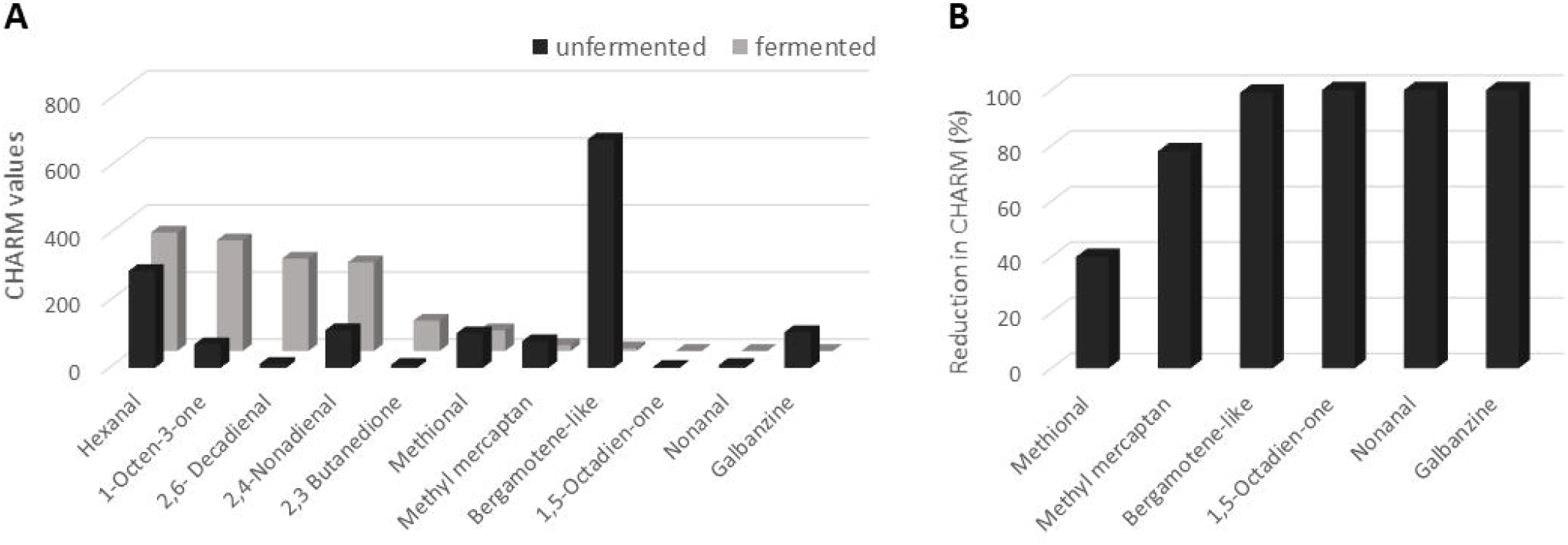
Quantification of odorants in fermented and unfermented protein blends. **A**) Levels of known volatiles compounds detected during CHARM analysis in fermented (back bars) and unfermented (gray bars) protein blends were expressed. **B)** Relative reduction of off-notes associated volatiles in fermented. Values were calculated as percentage of levels in unfermented and fermented protein blends.

## DISCUSSION

A major disadvantage of plant proteins is their comparatively lower nutritional quality relative to animal derived protein. Results of the ileal digestibility study demonstrated that PDCAAS was greater for the shiitake fermented protein compared with the unfermented protein, which indicates that the fermentation process may have changed the structure of the proteins and thereby made them more digestible. The observations that for both age groups, DIAAS values for the fermented protein was 23-24% greater than for the unfermented protein further indicates that fermentation increased the value of the proteins. Proteins with a DIAAS value between 75 and 100 are considered “good” sources of protein whereas proteins with a DIAAS >100 are considered “excellent” proteins (Leser, 2013); in this sense, the shiitake fermentation process transformed a good protein source into an excellent one for individuals older than 3 years. The relatively lower increase in PDCAAS versus DIAAS is likely because the fermentation of proteins in the hindgut equalizes the digestibility of protein between different sources even if the ileal digestibility of amino acids is different. The reason the PDCAAS values, regardless of protein and age group, were all greater than the DIAAS values is that although the same scoring pattern was used, the digestibility of crude protein, which is used in the calculation of PDCAAS values, was greater than the digestibility of the first limiting amino acid. However, because the digestibility of amino acids is more correctly estimated by the digestibility of the individual amino acids than by the digestibility of crude protein, the DIAAS values are more representative of the nutritional value of proteins than PDCAAS values.

Several factors might act synergistically to increase the digestibility of the protein blend during the fermentation. Fungi are known to secrete a wide variety of enzymes, including proteases. Shiitake secreted proteases might “pre-digest” the protein substrate before they reach the pig digestive system while the increased solubility of the fermented protein, especially at low pH, may partially account for the observed increased digestibility. Additionally, the level of the gastric enzymes’ inhibitor, phytate, was substantially reduced by the fungal fermentation process. It is very foreseeable that this lower phytate level also contributed to the observed increase in the pigs’ digestibility of the fermented protein blend. Genome searches of different publicly available shiitake genomes indicates that different strains contain at least 5 genes encoding predicted phytases in addition to additional genes encoding potential inositol polyphosphate phosphatases (https://mycocosm.jgi.doe.gov/mycocosm/home). Moreover, the presence of a signal peptide sequence at the N-terminus of most phytases, suggests that shiitake secretes a substantial amount of phytase that could act to degrade phytic acid during fermentation of pea and rice substrates, accounting for the approximately 46% reduction of phytate in the fermented blend. A substantial reduction in cysteine protease inhibition (papain) is observed during the fermentation process. Enzymatic microbial enzymatic activity during fermentation has also been shown to reduce gastric protein inhibitors from plant protein (Avilés-Gaxiola et al., 2018). On the other hand, the antinutrient papain inhibitor oryzacystatin-I is a protein itself, therefore the denaturation/degradation of this protein during sterilization process of the unfermented pea and rice protein blend could also partially contribute to the reduced enzyme inhibition in the fermented protein blend.

White-rot fungi, such as shiitake, secrete a cocktail of “lignin modifying enzymes” (LME) which catalyze the breakdown of lignin, an amorphous polymer present in the cell wall of plants and the main constituent of wood. LME are oxidizing enzymes and include manganese peroxidase (EC 1.11.1.13), lignin peroxidase (EC 1.11.1.14), versatile peroxidase (EC 1.11.1.16) and laccases (EC 1.10.3.2). Many LME have a low specificity and can oxidize a wide range of substrates with phenolic residues, beside lignin (Plácido & Capareda, 2015). For example, laccases oxidize a variety of phenolic substrates, performing one-electron oxidations, leading to crosslinking and polymerization (Eisenman et al., 2007) of the ring cleavage of aromatic compounds. Fungal laccases and tyrosinases oxidize phenolic residues in protein and carbohydrates present in wheat flour improving its baking properties (Selinheimo & Valtion teknillinen tutkimuskeskus, 2008). Moreover, shiitake laccases have been used to remove off-flavor notes from apple juice (Schroeder et al., 2008). Gene expression profiling (RNA-Seq) indicates that many laccase genes as well as other LMEs are expressed by shiitake, and a few are upregulated during the shiitake fermentation of pea and rice protein blend (data not shown). Therefore, it is very likely that shiitake LME oxidation of key phenolic residues in the protein blend accounts in part for the reduction/elimination of off-note compounds, resulting in improved organoleptic properties. Other mechanisms such as physical trapping of volatiles and thermal reactions during the sterilization and drying of the protein blends may also contribute to the changes in olfactory character. Further studies on the mode of action and combination of mechanisms responsible for the taste improving capacity of shiitake mycelium fermentation are ongoing.

### Conclusion

The benefits of fermentation on pea protein taste and aroma has been demonstrated by Schindler and colleagues^54^. However, to our knowledge, the work presented here is the first successful application of fungal fermentation for the improvement of plant-based protein concentration. The action of the fungal mycelium results in a reduction of compounds negatively impacting the organoleptic characteristics of plant proteins while improving the digestibility and reducing antinutrient contents. This pioneering work will most certainly serve as a basis for future application of mycelial fermentation to improve the quality of low-quality sources to meet the food standards associated with food ingredients.

---

**Supplementary Figure 1. Fold-change of detected oxylipins**. Levels of detected oxylipin derived compounds present in the fermented protein blend were normalized to value detected in the unfermented protein blend.

