## Supplementary table 1 for "Shiitake mycelium fermentation improves digestibility, nutritional value, flavor and functionality of plant proteins"

**Supplementary Table 1.** GC-FID, GC-O and CHARM analysis of volatile compounds in the fermented and unfermented protein blend samples.

| **CHARM**  **F** | **CHARM**  **UF** | **Odor Perception** | **Odorant** |
| --- | --- | --- | --- |
| 354 | 289 | Green | hexanal |
| 330 | 70.5 | Musty | 1-octen-3-one |
| 276 | 13 | Fatty | 2,6-decadienal |
| 264 | 113 | Green/fatty | 2,4-nonadienal |
| 90 | 9 | Buttery | 2,3-butanedione |
| 62 | 104 | Potato | methional |
| 31 | 1 | Malt | 3-methyl butanal |
| 20 | 6 | Fresh cut grass/ aldehyde | 2-nonenal |
| 18 | 80 | Mustard/cabbage | methanthiol |
| 8 | 682 | Earthy | bergamotene-like |
| 1 | 1 | Roasted/nutty | 2-methyl-3-furanthiol |
| 1 | 4 | Musty | 1,5-octadienone |
| 3 | 9 | Cucumber/aldehyde | nonanal |
| 1 | 107 | Beany | Galbazine |

Major hedonic compounds are described along with the presumed compound identity based on Kovacs RI values and the FlavorNet database (<https://www.flavornet.org/index.html>). **UF** unfermented protein blend; **F** fermented protein blend.
