## Supplementary table 2 for "Shiitake mycelium fermentation improves digestibility, nutritional value, flavor and functionality of plant proteins"

| **Supplementary table 2**. ADJUSTED Charm data file for Series A containing: | | | | | | |  |  |  |  |  |
| --- | --- | --- | --- | --- | --- | --- | --- | --- | --- | --- | --- |
| **RI** | **Charm** | | **FDV** | | **Perception** | **Odorant ID** | **MS** | **MS ion** | **CAS #** | **RI Match** | **Odor Match** |
| 612 | 26.6 | | 9 | | rotten cabbage | Methanethiol | Y | 47.48 | 74-93-1 | Y | Y |
| 638 | 21.2 | | 9 | | buttery | 2,3-Butanedione | Y | 48.60 | 431-03-8 | Y | Y |
| 666 | 19.3 | | 9 | | malt | 2-Methyl butanal | Y | 41.96 | 96-17-3 | Y | Y |
| 796 | 247.3 | | 81 | | green | Hexanal | Y | 72.82 | 66-25-1 | Y | Y |
| 899 | 4.4 | | 3 | | green | Heptanal | Y | 70.81 | 000111-71-7 | Y | Y |
| 978 | 17.7 | | 9 | | mushroom | 1-Octen-3-one | Y | 55.70 | 4312-99-6 | Y | Y |
| 984 | 1.4 | | 3 | | geranium | Octadienone | Y | 55.12 | 65767-22-8 | Y | Y |
| 1003 | 0.6 | | 1 | | green | Octanal | Y | 55.84 | 124-13-0 | Y | Y |
| 1058 | 1.7 | | 1 | | green-fatty | 2-Octenal | Y | 70.83 | 2548-87-0 | Y | Y |
| 1081 | 1 | | 1 | | nutty/roasted | unknown | N |  |  | N | N |
| 1090 | 0.5 | | 1 | | nutty/roasted | unknown | N |  |  | N | N |
| 1100 | 1.4 | | 1 | | paper | Nonanal | Y | 82.98 | 124-19-6 | Y | Y |
| 1147 | 6.8 | | 3 | | paper | 2-Nonenal | Y | 41.70 | 2463-53-8 | Y | Y |
| 1182 | 69.2 | | 27 | | bell pepper | methoxyisobutylpyrazine | Y | 124.15 | 24683-00-9 | Y | Y |
| 1202 | 0.8 | | 1 | | green-fatty | Decanal | Y | 82.11 | 112-31-2 | Y | Y |
| 1215 | 32.9 | | 9 | | green-fatty | E,Z-2,4-Nonadienal | Y | 81.13 | 25152-84-5 | Y | Y |
| 1246 | 4.7 | | 3 | | floral | 2-Decenal | Y | 55.12 | 3913-71-1 | Y | Y |
| 1319 | 2.4 | | 1 | | green-fatty | E,E-2,4-Decadienal | Y | 81.15 | 25152-83-4 | Y | Y |
| 1382 | 2.1 | | 1 | | metallic | Epoxy 2- decenal | N |  | 134454-31-2 | Y | Y |
