## Supplementary table 3 for "Shiitake mycelium fermentation improves digestibility, nutritional value, flavor and functionality of plant proteins"

| **Supplementary table 3.** ADJUSTED Charm data file for Series A containing | | | | | | | | | | |  | | |  | | |  | | |  | |  |
| --- | --- | --- | --- | --- | --- | --- | --- | --- | --- | --- | --- | --- | --- | --- | --- | --- | --- | --- | --- | --- | --- | --- |
| **RI** | **Charm** | **FDV** | | **Perception** | **Odorant ID** | | | | | **MS** | | | **MS Ion** | | | **CAS #** | | | **RI Match** | | **Odor Match** | |
| 615 | 5.5 | 3 | | rotted cabbage | | Methanethiol | | | | | Y | | | 47.48 | | | 74-93-1 | | | Y | | Y |
| 640 | 25.2 | 9 | | buttery | | 2,3-Butanedione | | | | | Y | | | 43.86 | | | 431-03-8 | | | Y | | Y |
| 667 | 112.6 | 27 | | malt | | 2-Methyl butanal | | | | | Y | | | 41.96 | | | 96-17-3 | | | Y | | Y |
| 799 | 53.1 | 27 | | green | | Hexanal | | | | | Y | | | 72.82 | | | 66-25-1 | | | Y | | Y |
| 979 | 55.3 | 27 | | mushroom | | 1-Octen-3-one | | | | | Y | | | 55.70 | | | 4312-99-6 | | | Y | | Y |
| 985 | 3.2 | 3 | | geranium | | Octadienone | | | | | Y | | | 55.12 | | | 65767-22-8 | | | Y | | Y |
| 1091 | 3.4 | 3 | | nutty/roasted | | unknown | | | | | N | | |  | | |  | | |  | |  |
| 1103 | 2.9 | 3 | | milky paper | | Nonanal | | | | | Y | | | 82.98 | | | 124-19-6 | | | Y | | Y |
| 1148 | 5.5 | 3 | | paper | | 2-Nonenal | | | | | Y | | | 41.70 | | | 2463-53-8 | | | Y | | Y |
| 1186 | 10.4 | 9 | | bell pepper | | Methoxyisobutylpyrazine | | | | | N | | | 124.15 | | | 24683-00-9 | | | Y | | Y |
| 1209 | 4.8 | 3 | | green-fatty | | unknown | | | | | N | | |  | | |  | | |  | |  |
| 1223 | 5.8 | 3 | | green-fatty | | unknown | | | | | N | | |  | | |  | | |  | |  |
| 1321 | 4.1 | 3 | | green-fatty | | EE 2,4-Decadienal | | | | | Y | | | 81.15 | | | 25152-83-4 | | | Y | | Y |
| 1386 | 3.1 | 3 | | metallic | | Epoxy 2- decenal | | | | | N | | | 68.81 | | | 134454-31-2 | | | Y | | Y |
