## Supplementary table 5 for "Shiitake mycelium fermentation improves digestibility, nutritional value, flavor and functionality of plant proteins"

| **Supplementary table 5.** Adjusted Charm data file for Series A containing: | | | | |  |  |  |  |  |
| --- | --- | --- | --- | --- | --- | --- | --- | --- | --- |
| MBOOST-RAW1X.D, MBOOST-RAW3X.D, MBOOST-RAW9X.D, MBOOST-RAW27X.D, MBOOST-RAW81X.D | | | | | | | | |  |
| **RI** | **Charm** | **FDV** | **Perception** | **Odorant ID** | **MS** | **MS Ion** | **Cas #** | **RI Match** | **Odor Match** |
| 613 | 9.8 | 3 | sour-rotted cabbage | Methanethiol | Y | 47.48 | 74-93-1 | Y | Y |
| 623 | 28.8 | 9 | green tea | Dimethyl sulfide | Y | 62.47 | 75-18-3 | Y | Y |
| 638 | 114.5 | 27 | buttery | 2,3-Butanedione | Y | 43.86 | 431-03-8 | Y | Y |
| 667 | 18.5 | 9 | malt | 2-Methyl butanal | Y | 41.96 | 96-17-3 | Y | Y |
| 688 | 93.7 | 27 | pungent | Pentanal | Y | 44.58 | 110-62-3 | Y | Y |
| 771 | 12.4 | 3 | green, apple | E-2-Pentenal | Y | 55.84 | 1576-87-0 | Y | Y |
| 798 | 5.2 | 3 | green | Hexanal | Y | 82.72 | 000066-25-1 | Y | Y |
| 899 | 0.9 | 1 | green | Heptanal | Y | 70.81 | 000111-71-7 | Y | Y |
| 906 | 24.2 | 9 | potato | Methional | Y | 48.10 | 3268-49-3 | Y | Y |
| 966 | 23.2 | 9 | rotted cabbage | Dimethy trisulfide | Y | 79.12 | 3658-80-8 | Y | Y |
| 977 | 350 | 81 | mushroom | 1-Octen-3-one | Y | 55,70 | 4312-99-6 | Y | Y |
| 982 | 1069.6 | 243 | geranium | Octadienone | Y | 55.12 | 65767-22-8 | Y | Y |
| 1056 | 4.4 | 3 | nutty/roasted | methyl ethyl pyrazine | T | 121.12 | 15707-23-0 | Y | Y |
| 1059 | 10.5 | 9 | green-fatty | 2-Octenal | Y | 70.83 | 2548-87-0 | Y | Y |
