## Supplementary figures and images for "Shiitake mycelium fermentation improves digestibility, nutritional value, flavor and functionality of plant proteins"

### Supplementary figure 1

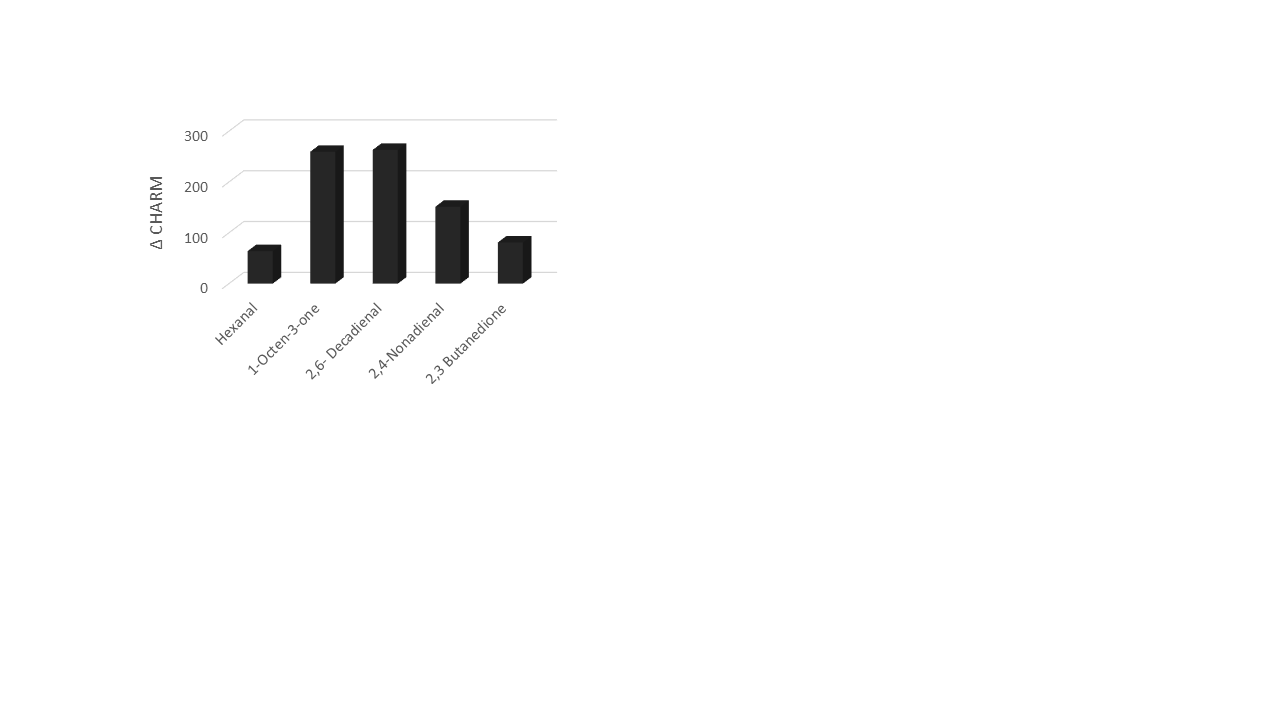
